# Alterations in Germinal Center Formation and B Cell Activation during Severe *Orientia tsutsugamushi* Infection in Mice

**DOI:** 10.1101/2023.01.12.523709

**Authors:** Casey Gonzales, Yuejin Liang, James Fisher, Galen Card, Jiaren Sun, Lynn Soong

**Affiliations:** Department of Pathology, University of Texas Medical Branch, Galveston, United States of America; Department of Microbiology and Immunology, University of Texas Medical Branch, Galveston, United States of America; Institute of Human Infections and Immunity, University of Texas Medical Branch, Galveston, United States of America; Sealy Institute for Vaccine Sciences, University of Texas Medical Branch, Galveston, United States of America

**Keywords:** Scrub typhus, humoral immunity, B cell, germinal center, splenic architecture, *Orientia tsutsugamushi*

## Abstract

Scrub typhus is a poorly studied but life-threatening disease caused by the intracellular bacterium *Orientia tsutsugamushi* (*Ot*). Cellular and humoral immunity in *Ot*-infected patients is not long-lasting, waning as early as one-year post-infection; however, its underlying mechanisms remain unclear. To date, no studies have examined germinal center (GC) or B cell responses in *Ot*-infected humans or experimental animals. This study was aimed at evaluating humoral immune responses at acute stages of severe *Ot* infection and possible mechanisms underlying B cell dysfunction. Following inoculation with *Ot* Karp, a clinically dominant strain known to cause lethal infection in C57BL/6 mice, we measured antigen-specific antibody titers, revealing IgG2c as the dominant isotype induced by infection. Splenic GC responses were evaluated by immunohistology, co-staining for B cells (B220), T cells (CD3), and GCs (GL-7). Organized GCs were evident at day 4 post-infection (D4), but they were nearly absent at D8, accompanied by scattered T cells throughout splenic tissues. Flow cytometry revealed comparable numbers of GC B cells and T follicular helper (Tfh) cells at D4 and D8, indicating that GC collapse was not due to excessive death of these cell subtypes at D8. B cell RNAseq analysis revealed significant differences in expression of genes associated with B cell adhesion and co-stimulation at D8 versus D4. The significant downregulation of *S1PR2* (a GC-specific adhesion gene) was most evident at D8, correlating with disrupted GC formation. Signaling pathway analysis uncovered downregulation of 71% of B cell activation genes at D8, suggesting attenuation of B cell activation during severe infection. This is the first study showing the disruption of B/T cell microenvironment and dysregulation of B cell responses during *Ot* infection, which may help understand the transient immunity associated with scrub typhus.

**Author Summary:** Scrub typhus is an understudied disease caused by the intracellular bacterium *O. tsutsugamushi*. A hallmark of scrub typhus is the unexplained, non-durable immunity after infection. While cellular immune responses are known to be important for controlling this infection, our understanding of B cell and GC responses remains limited. In this study, we examined B cell activation and GC responses using our recently established murine model of severe scrub typhus. We measured bacterial antigen-specific antibody titers and identified IgG2c, an IFN-γ-driven isotype, as the major IgG subtype. We also found that *O. tsutsugamushi* infection disrupted spleen morphology, exemplified by randomly dispersed T cells and lost GC structures. Transcriptomic analysis of purified splenic B cells demonstrated reduced expression of genes encoding critical adhesion and co-stimulation molecules, including GC-adhesion gene *S1PR2*, at severe stages of infection. Several humoral immune response pathways relevant to B cell receptor signaling, B cell activation and differentiation were significantly downregulated during infection. This study is the first report describing humoral immunity in a scrub typhus mouse model and provides detailed evidence that B cell and GC responses are impaired during acute infection.

## Introduction

Scrub typhus is a neglected but life-threatening infectious disease, which often presents as an acute undifferentiated febrile illness. It is caused by the obligately intracytosolic bacterium *Orientia tsutsugamushi* (*Ot*) that preferentially infects phagocytes and endothelial cells, replicates in the cytosol, and is released from host cells via a virus-like budding process [1]. It is estimated that one million individuals are infected each year, and nearly a third of the human population lives in endemic areas that are known as the “tsutsugamushi triangle” [2]. The recent emergence of scrub typhus in regions previously believed to be free of the disease has raised public health concerns [3–6]. *Ot* infects humans when larval *Leptotrombidium* mites feed on dermal tissue fluid. The bacteria then disseminate systemically to the major organs, leading to disease that ranges from mild to fatal illness. Progression of severe scrub typhus may manifest as hypotensive shock, acute respiratory distress, acute renal failure, meningoencephalitis, gastrointestinal bleeding, and coagulopathy, if diagnosis and appropriate antibiotic therapy are inadequate or delayed [3]. In experimental animal models, severe clinical outcomes and disease pathogenesis are positively correlated with excessive inflammation and type 1-skewed cytokine/chemokine production in multiple organs [7, 8]. In scrub typhus patients, proinflammatory cytokine and chemokine responses are characterized by upregulation of IFN-γ, TNF-α, IL-6, IL-8, IL-12p40, CXCL10, MCP-1, and MIP-1β, but downregulation of Th2-associated cytokines and chemokines [9–12]. Particularly, highly levels of TNF-α, IL-8, and IL-10 are positively associated with severe scrub typhus in humans [9, 11, 12]. While type 1 inflammation is essential for controlling bacterial replication, uncontrolled responses can lead to tissue injury and immune dysregulation [7–9, 13].

Clinical and epidemiological studies have suggested that the protective immunity elicited by *Ot* infection fails to be long-lasting, as reinfection can occur as early as one-year after primary infection in some individuals [14–19]. Scrub typhus patients display strong cellular immune responses, which are most critical for the control of *Ot* infection [20, 21]. Yet, cellular immunity wanes around one-year post-infection in humans, along with the decline of circulating antibodies within 1-2 years, following recovery from scrub typhus [16]. However, detailed evaluation of B cell responses or antigen-specific IgG isotype titers in scrub typhus patients is nearly absent. In experimentally infected non-human primates and mice, immune protection and circulating antibodies also seem to diminish quickly [15, 17]. Shirai *et al.* reported that passive transfer of anti-Gilliam and anti-Karp *Ot* sera protected 3 out of 5 tested mice against a lethal dose of Karp challenge [22]. Another report revealed that protection conferred by serum passive transfer was most effective at protecting naïve mice from lethal challenge with a homologous strain when the serum donor was infected multiple times [23]. These pioneer studies, however, were conducted a few decades ago with limited sample sizes and study depth. At present, there are no detailed animal studies discussing B cell responses in the context of *Ot* infection at the cellular/molecular level.

The germinal center (GC) reaction is critical for the development of durable humoral immunity against pathogens and is key to the adaptive immune response. Within GCs of secondary lymphoid organs, unique and dynamic cell-cell interactions occur between proliferating antigen-specific B cells, T follicular helper (Tfh) cells, and the specialized follicular dendritic cells. These interactions are designed to exert a selective pressure on B cells to increase the antigen specificity of their B cell receptors (BCRs) over time. This selection gauntlet only allows B cells with the most antigen-specific BCRs to differentiate into long-lasting memory B cells and long-lived plasma cells, respectively, which provide durable humoral immune protection after infection or immunization. Many bacteria, viruses, and parasites have developed unique mechanisms to subvert GC responses, interfering with the generation of long-lived humoral immunity. Some of the known mechanisms include altering B cell survival, eliciting excessive extrafollicular B cell responses [24, 25], inducing regulatory B cell differentiation (to shut down the immune responses) [26], or disrupting lymphoid tissue architecture [27, 28]. For example, i.p. inoculation of *Ehrlichia muris*, a closely related intracellular bacterium, elicited strong extrafollicular responses during infection, but delayed development of GCs [24, 29]. GC responses rely on precisely timed B and T cell migration as well as localization signals, orchestrated by tightly controlled chemokine gradients and upregulation of several receptors, including S1PR2 on GC cell subsets [30]. To initiate GC formation, antigen-activated B cells must migrate to the interfollicular region, where they interact with T cells. Following this interaction, a subset of B and T cells will upregulate CXCR5 and follow a CXCL13 chemokine gradient established by follicular dendritic cells that directs these cells to move toward the center of B cell follicles. Once localized at the center of the B cell follicle, these GC B cells, Tfh cells, and follicular dendritic cells will form a GC [31], in which B cells will then undergo the strict antigen-specific selection process described above. Several infections can alter lymphoid tissue organization and disrupt chemokine gradients that are essential for the initiation and maintenance of the GC reaction [32–37], ultimately leading to short-lived or attenuated humoral immunity. For example, Popescu *et al.* revealed that *E. muris* infection induced high levels of TNF-α, resulting in disrupted lymphoid tissue organization and GC B cell development in the spleen [33]. Similarly, pro-inflammatory mediators IFN-γ and TNF-α were implicated as drivers of splenic architecture modulation and inhibition of Tfh differentiation during severe malaria infection [35]. Yet, there are no reports detailing the GC reaction during *Ot* infection, leaving a critical knowledge gap regarding the humoral immune response to *Ot* and the transient immunity after *Ot* infection. Such knowledge deficit has also hampered scrub typhus vaccine development.

In this study, we examined B cell responses in detail by using our established severe scrub typhus model [38], via intravenous infection of C57BL/6 mice with *Ot* Karp strain. We used *Ot*-specific recombinant protein to confirm serum IgG2c as a dominant antibody isotype. Next, we used immunofluorescent staining and uncovered the disruption of GC structures in the spleens at severe stages (D8) of infection. Furthermore, by using RNAseq analysis of B cells purified from mock- and *Ot*-infected spleens, we identified downregulation of adhesion molecule and costimulatory genes, as well as impaired B cell activation and BCR signaling at late-stages of infection. This study provides the first lines of evidence, describing alteration in GC formation and humoral immune responses in mice during acute scrub typhus.

## Materials and Methods

### Mouse infection and ethics statement

Female C57BL/6J mice (#000664) were purchased from the Jackson Laboratory and maintained under specific pathogen-free conditions. Animals used were at 8-12 weeks of age, following protocols approved by the Institutional Animal Care and Use Committee (IACUC#1902006) at the University of Texas Medical Branch (UTMB) in Galveston, TX. All infection studies were performed in ABSL3 facilities in the Galveston National Laboratory. All procedures were approved by the Institutional Biosafety Committee, in accordance with Guidelines for Biosafety in Microbiological and Biomedical Laboratories. UTMB complies with the USDA Animal Welfare Act (Public Law 89-544), the Health Research Extension Act of 1985 (Public Law 99-158), the Public Health Service Policy on Humane Care and Use of Laboratory Animals, and the NAS Guide for the Care and Use of Laboratory Animals (ISBN-13). UTMB is registered as a Research Facility under the Animal Welfare Act and has current assurance on file with the Office of Laboratory Animal Welfare, in compliance with NIH policy. *Ot* Karp stocks were prepared in L929 cells. The inoculum infectivity titer was determined as in our previous report [38]. Mice were inoculated i.v. with a lethal dose (8 × 10^4^ FFU, 200 μl) of bacteria or PBS (mock) and monitored daily for weight loss, signs of disease, and survival. Serum and spleen tissue samples (3-5/group) were collected at D4 and D8 post-infection, followed by the subsequent immunological analysis. All experiments were performed with the same bacterial stock. Flow cytometry, IFA, and qRT-PCR data shown are representative of three independent repeats.

### Flow Cytometry

Single-cell suspensions were made by passing spleens through a 70-μm cell strainer in RPMI 1640 medium, and these were then treated with Red Blood Cell Lysis Buffer (Sigma-Aldrich). Cells were first blocked with FcγR blocker, followed by the staining of eFluor 506 Fixable Viability Dye and fluorochrome-labeled antibodies (Abs). The below Abs were purchased from either BioLegend or Thermo Fisher Scientific: PE-Cy7-anti-CD3, Alexa fluor700-anti-B220, Alexa fluor488-anti-GL-7, APC-anti-CD38. For the Tfh cell staining, cells were first incubated with the rat anti-mouse CXCR5 for 1 h, followed by the staining of secondary Ab biotin-conjugated AffiniPure Goat anti-rat (H+L) for 30 min, and tertiary Ab APC-streptavidin along with PE-Cy7-anti-CD3, PerCP-Cy5.5-anti-CD4, and FITC-anti-PD-1. Cells were fixed in 2% paraformaldehyde overnight at 4°C prior to analysis. Data were acquired on a BD LSR Fortessa in the UTMB Flow Cytometry Core and analyzed by using FlowJo software version 10.7.2 (BD Bioscience).

### Immunofluorescence Imaging

Spleen tissues were fixed in 4% paraformaldehyde for 24 h. Tissues were transferred into 20% sucrose/PBS for 24 h, followed by 30% sucrose/PBS for another 24 h. All fixations were performed at 4°C, and tissues were finally frozen in O.C.T. compound (Sakura Finetek). Frozen sections (7-μm) were blocked with 1% BSA and 0.3 M glycine in PBS for 30 min. They were then incubated with rat IgG2a anti-B220 (1:150), rat IgM anti-GL-7 (1:150), biotin anti-CD3 (7 μg/mL) Abs for 1 h at room temperature (BioLegend, clones RA3-6B2, GL-7, 17A2, respectively). Sections were then stained with secondary Abs, Alexa Fluor 594-conjugated mouse anti-rat IgG2a (1:50, clone MRG2a-83, BioLegend), Alexa Fluor 488-conjugated goat anti-rat IgM (1:50, clone A21212, Invitrogen), streptavidin cyanine 5 (5 μg/mL, BioLegend) for 1 h. All samples were stained with DAPI (Sigma-Aldrich). Staining with secondary antibodies and primary antibodies alone served as negative controls. For each section, at least 4-5 fields of each spleen section were imaged at the UTMB Optical Microscopy Core on a Zeiss LSM 880 confocal microscope (Carl Zeiss Microscopy LLC) equipped with ApoTome and Zen imaging software. The 405, 488, 561, and 633 excitation lasers under 63× oil immersion objective were used. Acquisition settings were identical among samples of different experimental groups and representative images are presented from each time point.

### NanoString Gene Expression Profiling

Mouse spleen samples were collected from mock (D0) and D8 infected mice, and stored in RNA*Later* (Ambion, Austin, TX) until extraction was performed. Total RNA was extracted by using the RNeasy Mini kit (Qiagen), and total RNA samples (3 mice/group, 200 ng/sample in ribonuclease-free water) were processed at the Baylor College of Medicine Genomic and RNA Expression Profiling Core (Houston, TX). Gene expression profiling was performed by using the nCounter platform and the Mouse Immunology Panel NanoString kit, which comprised 561 genes and 14 housekeeping genes (NanoString Technologies, Seattle, WA). Raw data were then normalized and analyzed by ROSALIND (https://rosalind.onramp.bio/), with a HyperScale architecture developed by ROSALIND, Inc (San Diego, CA). NanoString criteria was used for normalization, calculation of fold changes and calculation of p-values. ROSALIND follows the nCounter Advanced Analysis protocol of dividing counts within a lane by the geometric mean of the normalizer probes from the same lane. Housekeeping probes used for normalization were selected based on the geNorm algorithm as implemented in the NormqPCR R library [39]. P-value adjustment was performed using the Benjamini-Hochberg method of estimating false discovery rates. Clustering of genes for the final heatmap of differentially expressed genes was done by using the Partitioning Around Medoids method with the flexible procedures for clustering R library (Hennig, CRAN R package) that takes into consideration the direction and type of all signals on a pathway, the position, role, and type of every gene.

### Quantitative Reverse Transcriptase PCR (qRT-PCR)

Splenic tissues were collected in RNA*Later* (Ambion) and incubated at 4°C overnight for inactivation. Tissues were homogenized using metal beads in a BeadBlaster 24 Microtube Homogenizer (Benchmark Scientific) with RLT lysis buffer (Qiagen). Total RNA was extracted by using RNeasy Mini Kit (Qiagen), and the cDNA was synthesized utilizing iScript Reverse Transcription kit (Bio-Rad). cDNA was amplified in a 10 μL reaction mixture containing 5 μL of iTaq SYBR Green Supermix (Bio-Rad) and 0.5 μM each of gene-specific forward and reverse primer. qRT-PCR assays were performed on a CFX96 Touch Real-Time PCR Detection System (Bio-Rad), and PCR assays were denatured for 30 s at 95°C, followed by 40 cycles of 15 s at 95°C, and 60 s at 60°C. To check specificity of amplification, melt curve analysis was performed. Relative quantitation of mRNA expression was calculated utilizing the 2^-ΔΔCT^ method. Primers used in qRT-PCR analysis are listed in S1 Table.

### Quantitative PCR for tissue bacterial burden

Spleen samples were homogenized by using a FastPrep-24 homogenizer (MP Biomedical). DNA was then extracted by using a DNeasy Blood & Tissue Kit (Qiagen) and used for qPCR assays, as in our previous report [7]. The primers used were OtsuF630 (5’-AACTGATTTTATTCAAACTAATGCTGCT-3’) and OtsuR747 (5’-TATGCCTGAGTAAGATACGTGAATGGAATT-3’) (Integrated DNA Technologies). The copy number for the 47-kDa gene was determined by known concentrations of a control plasmid containing single-copy insert of the gene. Gene copy numbers were determined via serial dilution (10-fold) of the *Ot* Karp 47-kDa plasmid. Bacterial burdens were normalized to total nanogram (ng) of DNA per μL for each of the same samples. Data are expressed as the gene copy number of 47-kDa protein per ng of DNA.

### Enzyme-linked Immunosorbent Assay (ELISA)

Serum was separated from whole blood samples in blood separation tubes (BD Bioscience) by centrifuging at 9000 RCF for 2 min. Samples were frozen at −30°C, and once thawed, samples were inactivated with 0.09% sodium azide overnight at 4°C. For analysis of bacteria-specific Ab responses, 96-well plates were coated with 2 μg/mL of recombinant Karp type-specific antigen 56 (TSA56, generated by Genscript) in PBS and blocked with 1% BSA. Serum was diluted 1:2 until endpoint titers were determined. Detection was performed utilizing the following horseradish peroxidase-conjugated primary antibodies that were diluted 1:1000 in blocking buffer: goat anti-mouse IgG1 (A10551, Thermo Fisher Scientific), goat anti-mouse IgG2c (PA1-29288, Thermo Fisher Scientific), and goat anti-mouse IgM (1021-05, Southern Biotech). Visualizing reagent utilized was the 1-Step Ultra TMB ELISA Substrate Solution (Thermo Fisher Scientific). Optical density was measured on the BioTek Epoch microplate spectrophotometer.

### Splenic B cell isolation and RNAseq

B220^+^ B cells were isolated via negative selection from single-cell suspensions by using the mouse B Cell Isolation Kit (Miltenyi Biotec), following the manufacturer’s instructions. B cell samples were evaluated for purity (~97-98%, based on B220 expression assessed by flow cytometry), stored in RNA*Later* (Ambion), and used for RNA extraction (RNeasy Mini Kit, Qiagen). RNAseq analysis was performed by LC Sciences (Houston, TX) for RNA purity/quantity assessment via Bioanalyzer 2100 and RNA 6000 Nano LabChip Kit (Agilent), mRNA extraction, cDNA libraries construction, and sequencing via the Illumina Novaseq™ 6000. Raw data were normalized and analyzed by ROSALIND (https://rosalind.onramp.bio/), with a HyperScale architecture developed by ROSALIND (San Diego, CA). The Mus musculus genome build mm10 was used as a reference genome. Quality control analysis was done by using FastQC and RSeQC6 [40]. ROSALIND used HTseq4 [41] to quantify individual sample reads and DESeq2 R library [42] to normalize reads via Relative Log Expression. Enrichment was determined by hypergeometric distribution and was calculated relative to a background gene set that was relevant for the experiment. Several databases were referenced for gene set enrichment analysis including Pathway Interaction DB, REACTOME, KEGG, PANTHER, Gene Ontology Biological Process, and BioPlanet. Meta-analysis of D4 versus mock and D8 versus mock samples was performed to identify differentially expressed genes between D4 and D8 samples as well as significant signaling pathways.

### Statistical Analysis

Data were presented as mean ± standard error of mean (SEM). ELISA Ab titers, qRT-PCR, qPCR, and flow cytometry data were analyzed with one-way ANOVA and Tukey’s multiple comparisons test Post Hoc for comparisons between groups. All data were analyzed by using GraphPad Prism software. Statistically significant values are denoted as \**p* < 0.05, ** *p* < 0.01, *** *p* < 0.001, and \*\*\*\**p* < 0.0001, respectively.

## Results

### *O. tsutsugamushi* infection and antibody production

B cell research is nearly absent in the context of scrub typhus. Using a murine model of *Ot* infection, we first examined bacterium-specific antibody responses during lethal scrub typhus. We i.v infected C57BL/6 mice with a lethal dose of *Ot* and monitored body weight and survival daily. As shown in Fig 1A, body weight loss began at D4 and declined by about 25% until mice succumbed to infection from D9 to D11. At D8, one day prior to death, we found substantial and statistically significant levels of splenic bacterial burden (Fig 1B). To assess humoral immune responses during infection, we measured the titers of serum *Ot*-specific antibody against *Ot* type-specific antigen 56 (TSA56), a well-characterized, immunogenic outer membrane protein [43, 44].

**Fig 1.**
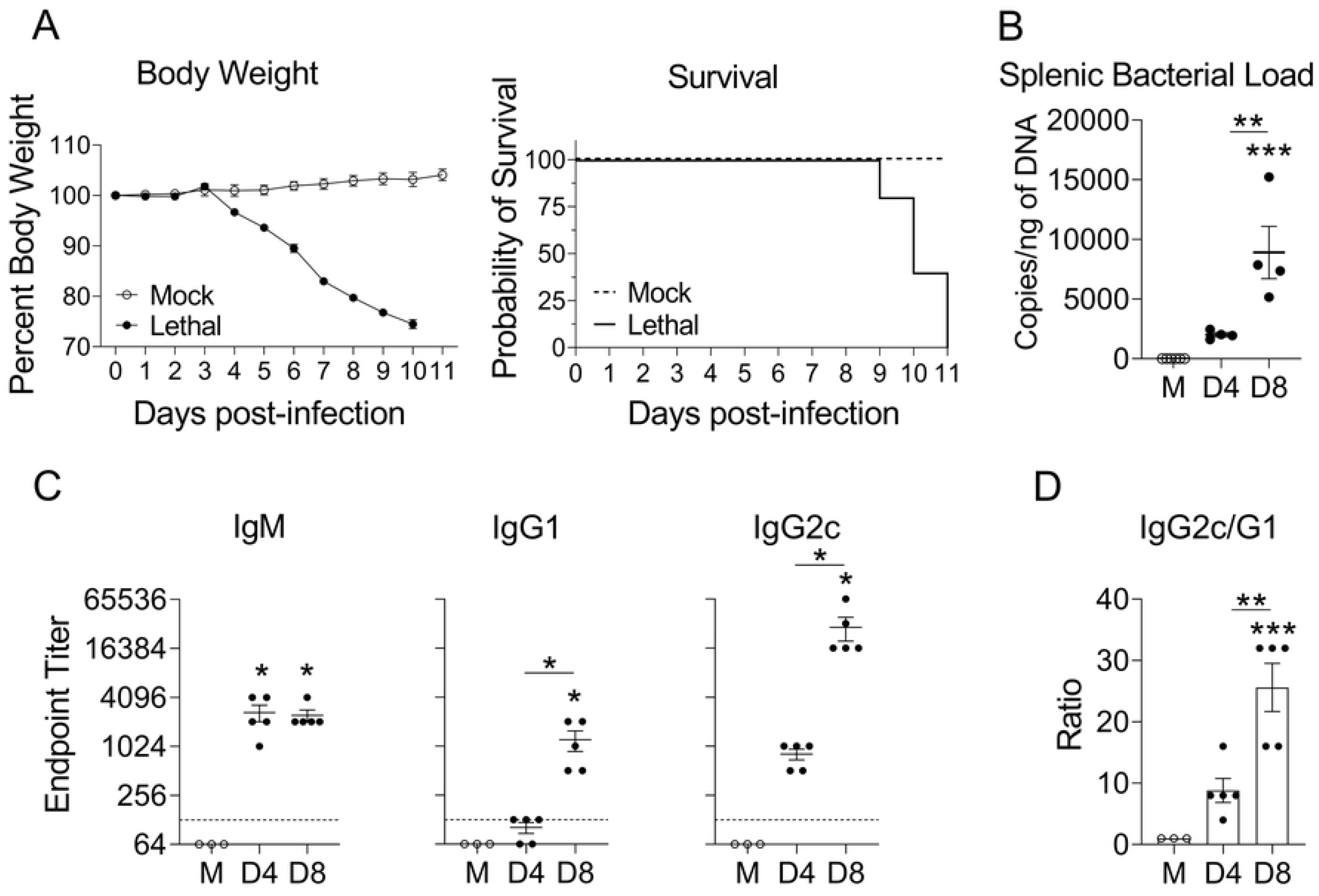
Disease progression and antigen-specific antibody titers in mice with lethal *O. tsutsugamushi* infection. A) C57BL/6 mice (n = 5) were i.v. infected with *O. tsutsugamushi* Karp strain (8 × 10^4^ FFU) and monitored daily for survival and body weight. PBS was used as a mock control. B) Mice (n = 3-4) were euthanized at D0 (Mock), D4 and D8, and spleens were harvested for bacterial load measurement by qPCR. C) Serum antibody titers specific to recombinant TSA56 were measured in infected mice (n = 5) and mock-infected mice (n = 3) via indirect ELISA assay; data were shown as endpoint titers of IgM, IgG1, and IgG2c, respectively. D) The IgG2C/IgG1 ratio was calculated. Data are shown as mean ± SEM from single experiments and are representative of two independent experiments with similar trends. For statistical analysis, one-way ANOVA was used with a Tukey’s multiple comparisons test. Asterisks without underlines were representative of comparison to mock. *,*p* <0.05; **,*p* <0.01; ***,*p* <0.001.

While IgM titers (1:2,048) were significantly increased in D4 and D8 samples, only D8 samples showed a statistically significant increase in IgG1 and IgG2c titers (Fig 1C). Of note, endpoint titers for IgG2c (1:32,768) at D8 were significantly higher than those of IgG1 (1:1,024), reaching an average IgG2c/IgG1 ratio of 25 (Fig 1D). These results reveal that the IgG response was predominantly composed of the IgG2c isotype, a finding that was reminiscent of that in *E. muris*-infected C57BL/6 mice [45]. Our finding of high IgG2c/IgG1 ratio suggested a type 1-skewed response *in vivo*, which was in accordance with our previous reports for strong Th1/IFN-γ, but weak Th2/IL-4-related, immune responses during severe *Ot* infection [7, 38, 46, 47].

### Dysregulated B and T cell homing gene expression during *O. tsutsugamushi* infection

To investigate the splenic B cell responses during *Ot* infection, we used infected spleens to analyze transcript levels of key molecules associated with antibody isotype switching and B cell recruitment and migration by qRT-PCR. IL-6, a cytokine involved in B cell proliferation and IgM isotype switching [48], initially increased in expression at D4, but dropped by approximately 66% at D8 (Fig 2A). This finding was consistent with our ELISA results demonstrating increased IgM endpoint titers at D4 and a sustained level at D8 (Fig 1C). We also assessed the expression of *IL-4* and *IFN-γ,* which are known to induce IgG1 and IgG2c isotype switching, respectively [48, 49]. At D8, we found that *IL-4* expression was reduced by 42%, while *IFN-γ* expression levels were nearly 22 times higher relative to mock-infected mice (Fig 2A). From D4 to D8, *IFN-γ/IL-4* ratio increased almost 250%, indicating a type 1-skewed immunity, especially at severe stages of infection. Since IL-21 shares the common receptor γ-chain with IL-4, contributing to isotype switching to IgG1 and initiation and maintenance of GC B cell responses [50–53], we examined *IL-21* transcripts and found a significant repression at both D4 and D8, as compared to mock-infected mice (Fig 2A, *p* < 0.001), indicating impaired IgG1 responses during infection. To gain insight into the homing signals of B and T cells, we measured chemokine receptor transcript levels and found 77% and 83% reduction of *CXCR5* and *CCR7* at D8 relative to mock-infected mice, respectively (Fig 2A).

**Fig 2.**
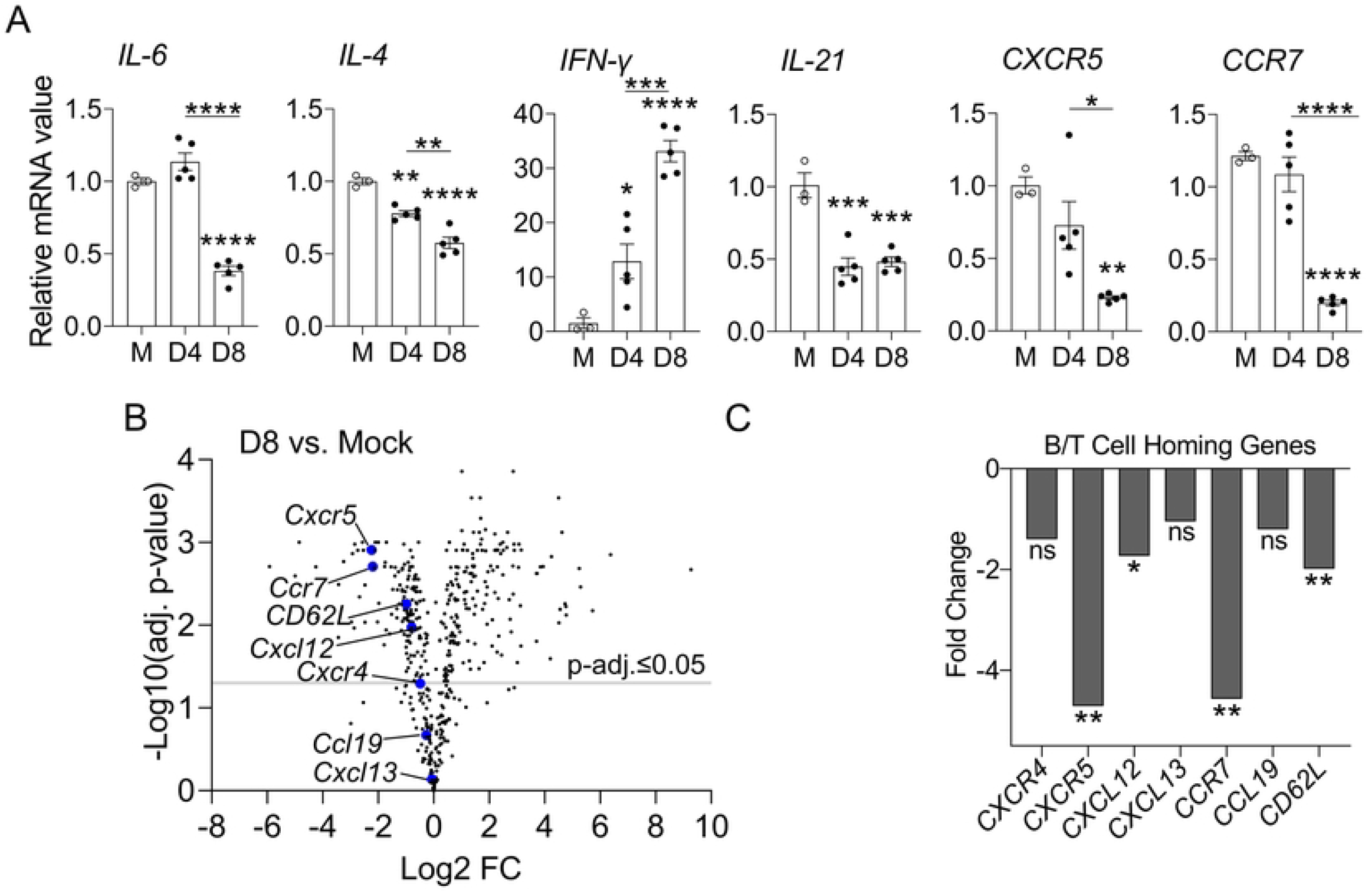
Splenic transcriptional profiles of antibody isotype-switch- and B/T cell recruitment-related genes during infection. A) Mice (n = 3-5) were infected, as described in Figure 1. Spleen tissues were harvested at indicated time-points. The transcript levels of *IL-6, IL-4, IFN-γ, IL-21, CXCR5* and *CCR7* were analyzed by qRT-PCR. A one-way ANOVA, with a Tukey’s multiple comparisons test was used for data analysis. Data are shown as mean ± SEM from single experiments and are representative of two independent experiments with similar trends. B) RNA samples extracted from the spleens of mock-infected and D8-infected mice (n = 3) were analyzed by Nanostring technology using the nCounter Immunology Panel Kit. Differential gene expression was analyzed via ROSALIND (https://rosalind.onramp.bio/). The key molecules for B/T cell homing were highlighted using blue circles, C) and their Log2 fold change (Log2 FC) changes as well as adjusted *p*-values were plotted. The statistical analysis was assessed via Benjamini-Hochberg test. *, *p* <0.05; **, *p* <0.01; ***, *p* <0.001; ****, *p* <0.0001. Asterisks without underlines were representative of comparison to mock.

To further explore alterations in humoral immunity, we examined B/T cell homing molecules through differential expression analysis between mock and D8 samples via NanoString. All genes encoding chemokines and their respective signaling receptors related with splenic B/T cell homing were downregulated at D8 (Fig 2B). Notably, *CXCR5* and *CXCL12* (chemokines critical for B cell homing and migration) were significantly downregulated, with a fold change of −4.71- and −1.73, respectively. In addition, T cell homing genes *CCR7* and *CD62L* were also reduced by −4.57- and −1.99-fold changes, respectively (Fig 2C). Overall, our results indicated high levels of type 1 immunity (e.g., IFN-γ) and suppressed type 2 responses (e.g., IL-4) in the spleen, reaffirming findings in our previous reports [7, 38, 46, 47]. Moreover, the downregulation of key splenic B/T cell homing genes may implicate poor B and T cell localization in follicles and T cell zones, especially during severe stages of infection.

### Abrogation of splenic architecture and disordered GC structures at severe stages of infection

Available reports involving either human scrub typhus patients or experimentally infected animals have indicated that antibodies are not long-lasting and can wane as early as one-year post-infection [14–16]. Since GCs are the site of developing memory B and long-lived plasma cells, we analyzed the formation of GCs via immunostaining of spleen sections for B cells (B220^+^), T cells (CD3^+^), and GCs (GL7^+^). Using confocal microscopy, we detected baseline GCs in mock spleens and found unique architectural trends of T cell zones and GCs during the infection (Fig 3). In D4 spleens, the GC expansion was evident, judged by distinct GC clusters formed within the B cell follicles and expanded T cell zones. However, D8 spleens displayed notable disorganization of lymphoid structures. Although B, T, and GC B cells were readily detectable at D8, we found that only B cells were maintained in discernable follicles, while GC B cells seemed to be dispersed randomly throughout the spleen tissue and failed to form distinguishable GCs. T cells were also scattered throughout the spleen at D8, with few organized T cell zones remaining. Collectively, these immunohistology results suggested impaired GC formation at severe stages of infection.

**Fig 3.**
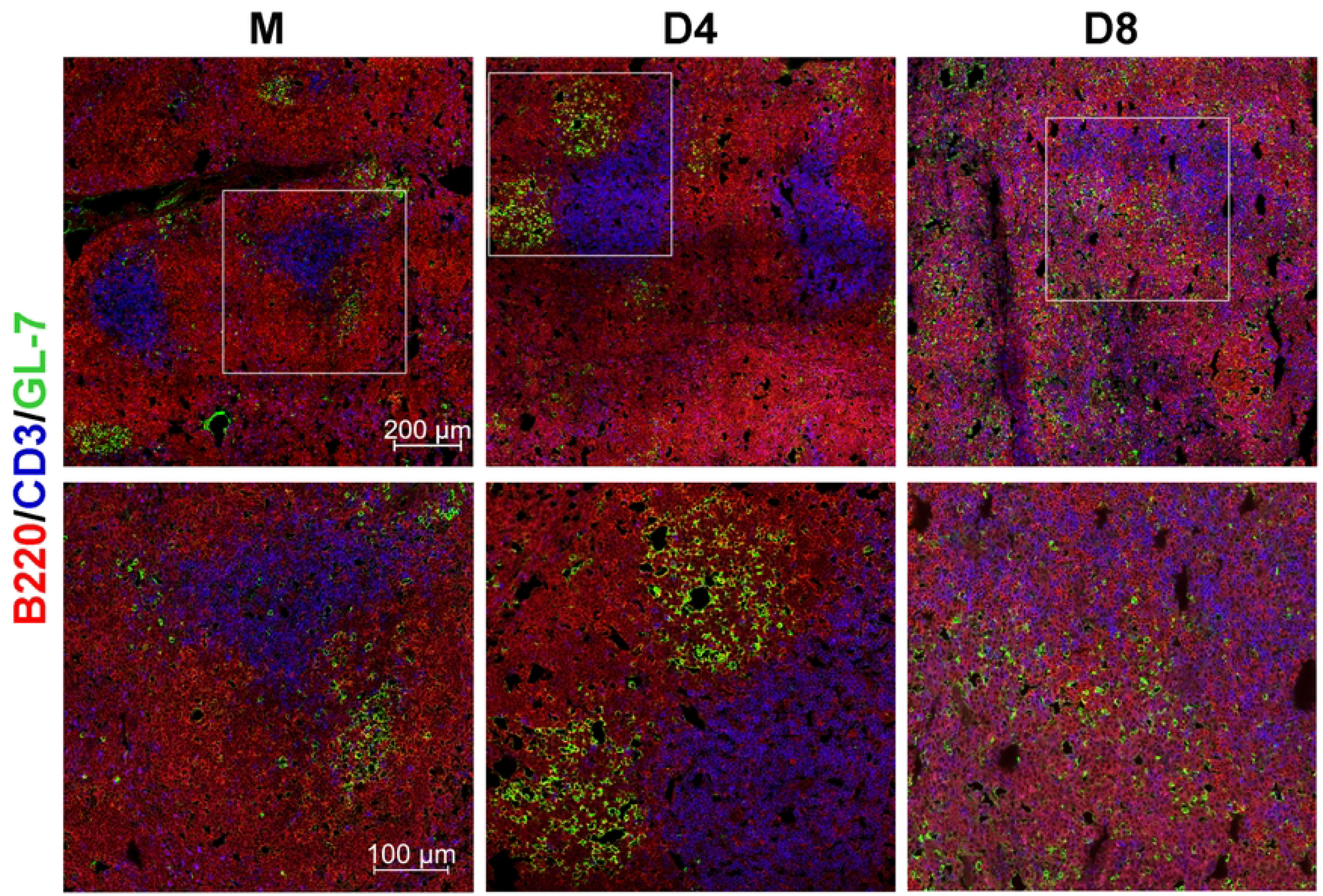
Disrupted GC formation and splenic architecture at the severe stage of infection. Spleen tissues were harvested from mock and infected mice, as described in Figure 1. Frozen sections of spleen (7 μm) were prepared and co-stained for B220 (total B cells, red), GL-7 (GC B cells, green), and CD3 (T cells, blue). Images were captured using confocal microscopy and are shown at low magnification (scale bar, 200 μm). Selected regions (gray boxes) are shown in high magnification (scale bar, 100 μm). Representative images were acquired from at least three independent mouse infection experiments with similar results.

### Impaired B cell responses in severe *O. tsutsugamushi* infection

GC responses are critical to the development of long-lived humoral immunity. To confirm and expand our findings of altered GC responses at late disease stages, we analyzed several splenic cell populations by multi-color flow cytometry (Fig 4A). B cell numbers significantly increased at D4 but declined to the level of mock samples at D8, indicating a selective impairment in B cell responses at the peak of infection (Fig 4B). While we revealed evidence of scattered GC B cells at D8 (Fig 3), we found that this disorganization did not correspond with a reduction in GC B cell numbers, as the number of GL-7^+^ B cells increased by nearly 3-fold at D4 and was sustained at D8 (Fig 4B). We next evaluated CD4^+^ T and Tfh cells (Fig 4B), as they are known to provide help to both B cells and GC B cells. The total number of CD4^+^ T cells was significantly increased at D4 but was reduced to nearly mock levels at D8 (Fig 4B). Total Tfh cell numbers were significantly increased by approximately 10-fold at D4, and these numbers were not significantly changed at D8 (Fig 4D). Therefore, the loss of CD4^+^ T and B cell numbers may lead to poor humoral immunity during acute *Ot* infection.

**Fig 4.**
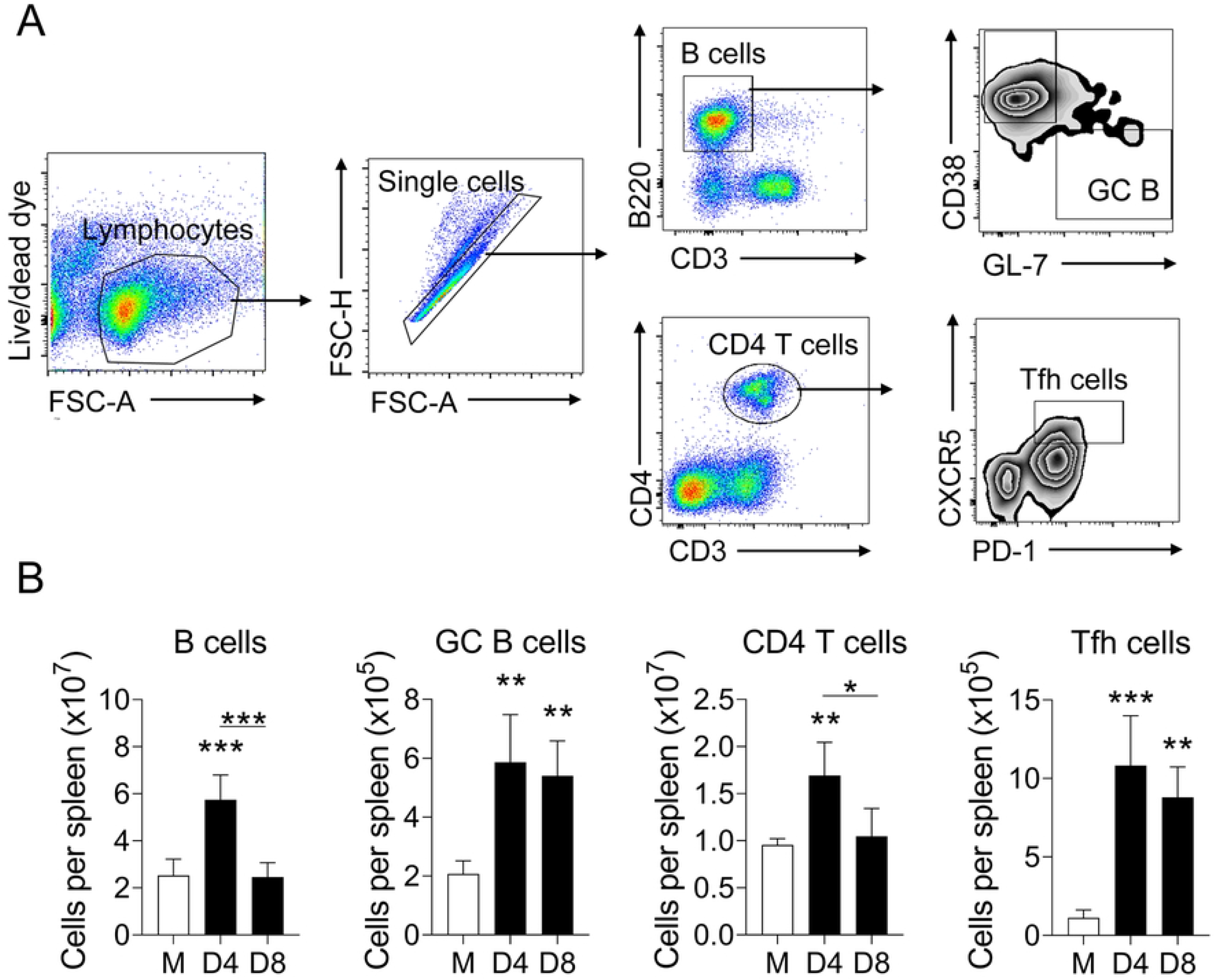
Splenic B and T cell subsets in *O. tsutsugamushi-infected* mice. Spleen tissues were harvested from mock and infected mice (n = 4), as described in Figure 1. Single-cell suspensions were prepared from spleens and stained for indicated cell surface markers, followed by flow cytometric analysis. Results were presented as A) gating strategy and B) absolute numbers of different cell populations. Data are shown as mean ± SEM from single experiments and are representative of three independent experiments with similar trends. For statistical analysis, a one-way ANOVA was used with a Tukey’s multiple comparisons test. *,*p* <0.05; **,*p* <0.01; ***,*p* <0.001; ****,*p* <0.0001. Asterisks without underlines were representative of comparison to mock.

### *O. tsutsugamushi* infection modulated B cell adhesion and co-stimulatory genes

It is known that the process of GC formation and maintenance involves carefully timed upregulation of several genes associated with B cell migration [30, 54, 55], and that inhibition of this process can result in the loss of GCs [56, 57]. To explore the mechanisms underlying the disorganization of splenic architecture during infection (Fig 3), we performed RNAseq on purified splenic B cells, and performed a meta-analysis of D4 and D8 samples, normalized to mocks. Overall, there were 2286 (1207 upregulated, 1079 downregulated) differentially expressed genes at D4, and 3496 (1988 upregulated, 1508 downregulated) differentially expressed genes at D8 (S1 Fig). We detected nine significant genes associated with B cell localization, five of which were downregulated on both D4 and D8, including *S1PR1, ITGA4, ITGA6, GPR183,* and *CCR7* (Fig 5A). Three of these genes, *S1PR1, GPR183,* and *CCR7,* showed a trend of continuous downregulation throughout infection, with fold-changes of −1.68, −2.30, and −1.64 at D8, respectively (Fig 5A). We also analyzed the normalized expression of *S1PR2*, which is known to be critical for confinement of GC B cells within GCs [56]. We found that *S1PR2* expression was significantly upregulated at D4 (2.18 fold-change) but dropped at D8 (to −1.30 fold-change, Fig 5B). Regarding GPR183 (also known as EBI2), a receptor known to be crucial for B cell migration to T cell zones [57–59], we found a significant reduction in *GPR183* expression at D4 (−1.34 fold change), which continued decline to −2.30 fold-change at D8 (Fig 5B). Together, these results suggest that the downregulation of genes associated with adhesion molecules, especially genes encoding the critical receptors *S1PR2* and *GPR183*, may contribute to the loss of GCs and disrupt of B/T cell interaction at the severe stage of infection.

**Fig 5.**
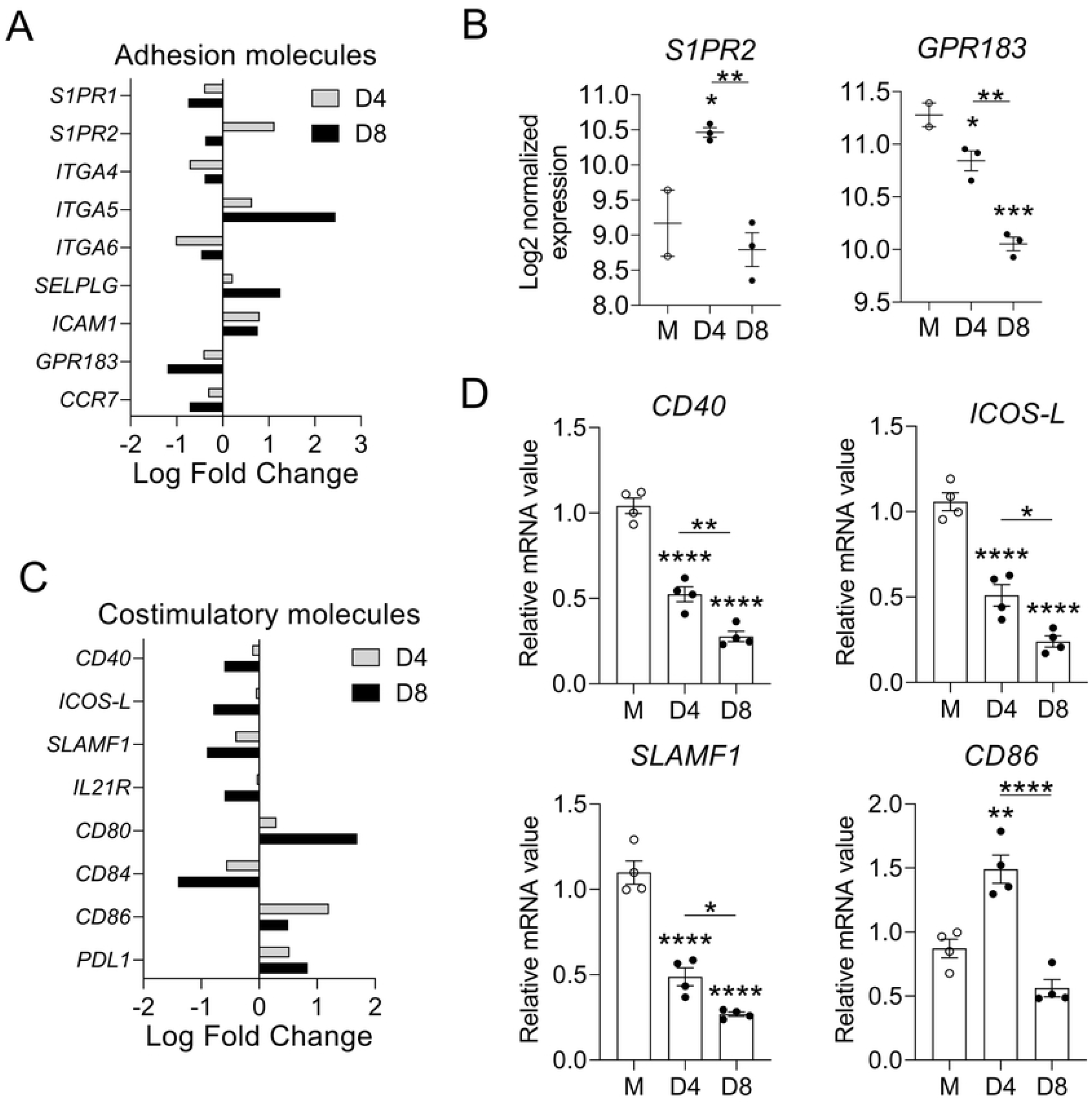
Alterations in genes encoding B cell-homing molecules and T/B cell interaction markers. A-C) Mice were infected, as described in Figure 1; splenic B cells were purified from mock samples (n = 2) and at D4 (n =3) and D8 (n =3) for RNAseq analyses. Data were normalized via Relative Log Expression (RLE) using DESeq2 R library. A, C) Meta-analyses between D4 and D8 samples were employed by ROSALIND with Log fold change of adhesion and costimulatory molecule genes shown, respectively. B) Selected genes for B cell adhesion molecules are shown in Log2 normalized expression. D) The transcript levels of B/T cell interaction markers on purified B cells were analyzed by qRT-PCR. For statistical analysis in panel B and D, the one-way ANOVA was used with a Tukey’s multiple comparisons test. * *p* <0.05; ** *p* <0.01; *** *p* <0.001; **** *p* <0.0001. Asterisks without underlines were representative of comparison to mock.

Strong and long-lasting GC responses are dependent on the second signal of B cell activation, including co-stimulation of B cells by Tfh cells [54, 55, 60]. Therefore, we next investigated the expression of genes encoding costimulatory molecules on B cells. Our meta-analyses indicated that five out of the eight significantly identified genes (*CD40, ICOS-L, SLAMF1, CD84, IL-21R*) were downregulated continuously during infection (Fig 5C). While expression of *CD80* (*B7-1*) and *PDL1* was continuously upregulated throughout infection, *CD86* (*B7-2*) expression increased initially at D4 but dropped by 0.41-fold at D8 (Fig 5C). We also confirmed some of our RNAseq findings by performing qRT-PCR (Fig 5D). Among molecules that have known roles in GC B cell survival (*CD40*, *ICOS-L*, *SLAMF1*, *CD84*, and *IL-21R*) [54, 61], their downregulation during *Ot* infection was highly significant and time dependent. Overall, our findings suggested a profound failure in upregulating most costimulatory molecules necessary for B/T cell interactions during *Ot* infection, especially those that are key signals for GC B cells.

### Diminished B cell activation at severe stages of *O. tsutsugamushi* infection

Since many bacterial species have evolved to manipulate signaling within B cells as an immune evasion tactic [26], including the enhancement or impairment of B cell activation, we then compared differentially expressed B cell-activation genes identified by B cell RNAseq. Using PANTHER and Biological Process databases, we searched for significant genes associated with B cell activation within the differentially expressed genes at D8 normalized to mocks. We found a total of 46 genes related to B cell activation, which demonstrated an adjusted *p*-value less than 0.05 and fold change greater than 1.5 or less than −1.5 (Fig 6A). Among them, 33/46 (71.7%) were downregulated, and only 13/46 (28.3%) genes were upregulated at D8 (Fig 6A). This observation was in sharp contrast to the expression of B cell activation genes at D4, of which 15/26 (57.7%) genes were upregulated (S2 Fig). At D4, among 26 differentially expressed genes related to B cell activation, 15 genes were upregulated and 11 genes were downregulated (S2 Fig).

**Fig 6.**
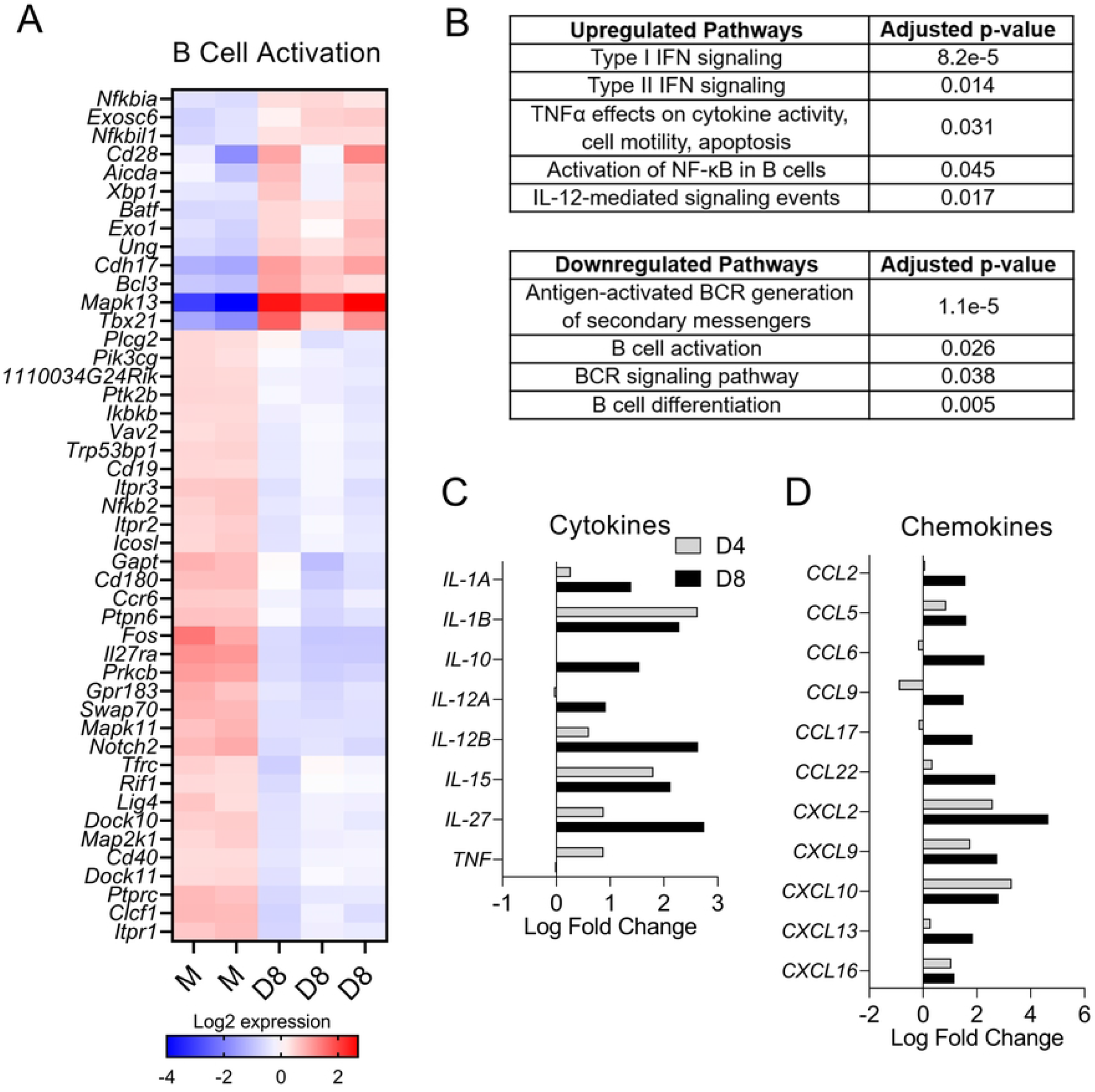
Impaired B cell activation in mice following *O. tsutsugamushi* infection. RNAseq data of splenic B cells were generated, as described in Figure 5. A) Differentially expressed genes for B cell activation (PANTHER and Gene Ontology Biological Process databases) from mock and D8 samples were identified by ROSALIND analysis and listed in the heatmap as subtracted normalized Log2 expression values. B) Meta-analysis between D4 and D8 samples identified a list of up- and down-regulated pathways (Pathway Interaction DB, REACTOME, PANTHER, Gene ontology Biological Process, and BioPlanet databases), and C-D) Log fold changes of differentially expressed C) cytokine and D) chemokine genes were shown.

Further analysis of genes associated with signaling pathways revealed a unique set of upregulated genes relating to a type 1-skewed response. We have reported the upregulation of IFN-γ and TNF-α in murine spleens during *Ot* infection [7]. As shown in Fig 6B, we found that B cells significantly upregulated genes relevant to signaling pathways mediated by type I IFN, type II IFN, TNF-α, and IL-12. In contrast, B cell activation genes and pathways upstream of B cell activation were significantly downregulated, including BCR signaling (S3 Fig) and secondary messengers for the BCR signaling. Likewise, B cell differentiation genes (*BCL10, CD72, FOS, PTPRC, SH3BP5, PTPN6*, etc.) were also downregulated throughout the infection, correlating with the drop of B cell numbers at D8 (Fig 4). Collectively, results in Figs 6A and B demonstrate an impairment in the humoral immune response to *Ot* infection, likely due to reduced expression of genes associated with B cell activation, BCR signaling, and other critical pathways to the B cell immunity.

There is currently a lack of understanding as to what key cytokines or chemokines B cells may produce in response to *Ot* infection. As shown in Fig 6C, we found the upregulation of *IL-12A*, *IL-12B*, and *TNF*, suggesting a possible involvement of B cells in shaping the splenic type-1 cytokine environment in *Ot* infection. We and others have previously reported upregulated IL-10 in *Ot*-infected mice and humans [7, 62-65]. Of note, we found an increase in B cell *IL-10* transcript levels from 1.01 fold-change at D4 to 2.92 fold-change at D8 (Fig 6C). To gain insight into the role of B cells in cell recruitment, we analyzed the chemokine gene expression in B cells and found the top upregulated chemokine genes included *CXCL2, CXCL9, CXCL10*, and *CCL22* (Fig 6D). Thus, B cells could play a role in recruiting inflammatory cells such as neutrophils (*CXCL2*) and CD8^+^ T/NK cells (*CXCL9*, *CXCL10*), but also anti-inflammatory T cells, such as Treg cells (*CCL22*). Since B cells also expressed *CCL2, CCL5,* and *CCL6,* they may play a role in monocyte and macrophage trafficking to the spleen as well (Fig 6D). Together, these results suggested a possible role for B cells to promote both type 1 immune responses and immunoregulatory responses via upregulation of cytokines and chemokines.

## Discussion

The emergence of *Orientia* infections into different geographic areas outside of its endemic distribution calls for vaccine-based controls. Despite the public health threat posed by this disease, detailed studies of adaptive immunity, especially those regarding humoral immunity, are scarce. Humoral immunity can contribute to host protection against *Ot* infection, as both hyperimmune serum and combined anti-Karp and anti-Gilliam sera can provide partial protection against lethal *Ot* challenge in mice [22, 23]. Yet, a scrub typhus vaccine is not available, partially due to a poor understanding of pathogenic mechanisms and host protective immunity. In this study, we used our established mouse model and performed tissue and cellular level transcriptional and immunological analyses to reveal dynamic patterns of B cell responses to *Ot* infection. We have provided the first lines of evidence that help understand the mechanisms underlying poor humoral immunity against *Ot* infection in mice.

First, we revealed TSA56-specific IgG2c-skewed antibody responses and a significantly high IgG2c/IgG1 ratio during infection, which were in agreement with, but more advanced than our previous studies that employed whole *Ot* bacterial lysates as the coated antigens for ELISA [7, 46]. These antibody titer trends were consistent with cytokine levels in the spleen at D8 (Fig 2A), namely reduced expression of *IL-6* and *IL-21* (IgM-switching) and *IL-4* (IgG1-switching), but a significant increase of *IFN-γ* (IgG2c-switching). B cell transcriptomics further revealed a selective upregulation of type 1 cytokine-related signaling pathways (type I IFN, type II IFN, TNF-α, IL-12, etc., Fig 6B), as well as upregulated type 1 cytokine-encoding genes (*IL-12A, IL-12B, TNF,* etc., Fig 6C), suggesting a possible contribution of B cells in promoting the type 1 immune environment. These findings support and extend our previous reports of Th1-skewed, but Th2-suppressed, immune responses during severe scrub typhus [7].

Second, we provided evidence of abrogated splenic architecture and disrupted GC structures at D8 of *Ot* infection. In other infectious disease models such as the self-limited *E. muris* infection in C57BL/6 mice [24, 33], the disorganization of secondary lymphoid organs and suppressed GC B cell responses are known to compromise humoral immunity and memory [34–36]. Using immunostaining (Fig 3), we found that D4 spleen sections initially contained expanded GCs and T cell zones, with distinct B cell follicles and T cell zones. However, D8 samples showed severe disruption of splenic architecture, characterized by scattered T cells with indistinct T cell zones. In contrast to the well-formed GC structures at D4, GC B cells were dispersed throughout the spleen, and no discernable GCs were maintained at D8. Since GCs rely on highly organized structure to properly initiate and sustain the GC reaction [55], the loss of GC structures indicates a lack of functional GC response at late-stage *Ot* infection.

We speculated that the distorted spleen morphology at severe stages of *Ot* infection may be partially attributed to the downregulation of B and T cell homing molecules (Figs 2B and 2C) since these molecules are critical to maintain normal architecture [28] and initiate GC responses [30]. B cell transcriptomics indicated significant downregulation of several adhesion molecules during infection, including *S1PR1, S1PR2, ITGA4, ITGA6, GPR183*, and *CCR7* (Fig 5A). Notably, the expression of *S1PR2*, responsible for confinement of GC B cells to GCs, was transiently upregulated at D4, but its expression dropped below the mock levels at D8 (Fig 5B), when GCs were lost. Therefore, reduced expression of GC adhesion gene *S1PR2* may play a role in the aberration of GC structures. The downregulation of *CCR7*, which drives B cell migration to the B/T zone, may also contribute to this dysregulated humoral immune response by reducing cognate interactions between B and T cells, as CCR7 is required for B cells to migrate to the B-T boundary. It has been reported that altered splenic morphology observed during *E. muris* infection was mediated partly by TNF-α, as infected TNF-α-deficient mice showed increased GC B cell numbers, clearly formed GCs, and distinct B and T cell zones as compared to WT-infected mice [33]. On the other hand, proinflammatory cytokines (TNF-α and IFN-γ) were implicated in GC and splenic architecture disruption, as their neutralization increased GC B cell number and improved splenic morphology and GC structures [35]. Since *Ot* infection elicits strong type 1 immune responses [7, 38, 47], these proinflammatory cytokines may contribute to the observed aberrant splenic architecture, and this direction is warranted for future study.

Third, we found impairment of several pathways critical to the humoral immune response to infection, including B cell activation and BCR signaling. Our RNAseq data exhibited 46 B cell activation-associated genes that were significantly changed at D8; among them, over 70% of genes were downregulated (Fig 6A). Moreover, other crucial signaling pathways in B cells were downregulated from D4 to D8, including BCR signaling, antigen-activated BCR generation of secondary messengers, and B cell differentiation (Fig 6B). Negative modulation of B cell activation has been observed with other bacterial infections [26]. For example, experimental infection of primary B cells with *Yersinia pseudotuberculosis* revealed diminished B cell activation due to the virulence factor yopH-mediated impairment of downstream BCR signaling events [66]. Thus, investigation of the *Ot*-specific negative modulation of BCR signaling and activation is necessary.

To explain the attenuation of B cell activation during infection, we examined the second signal of B cell activation. For a B cell to become activated, two signals are required: engagement of the BCR by antigen and co-stimulation by Th cells [67, 68]. To investigate this question, we examined the expression of genes encoding B cell costimulatory molecules and found that the majority were downregulated at D8 (Fig 5C). Some of the most important interactions for GC formation between B and Th cells are mediated through the costimulatory molecules CD40, ICOS-L, and SLAM proteins on B cells, which bind to their corresponding receptors (CD40L, ICOS, and SAP) on T cells, leading to costimulatory signal transduction and B cell activation [55, 60, 69, 70]. Our data showed that the genes associated with these key costimulatory molecules were significantly downregulated, indicating the dampened B cell costimulatory signaling pathways in *Ot* infection (Figs 5C and 5D). Together, these findings reveal that B cell activation is attenuated during *Ot* infection, and this phenomenon may be mediated by the inhibition of both the first (BCR signaling) and second (co-stimulation by Th cells) signals of B cell activation.

Our study demonstrated the dysregulated B cell responses in severe *Ot* infection; however, due to the lethality of the high dose i.v. murine model, only two time-points of infection were explored in this study. Future studies involving lower inoculation doses, more time points, and/or subclinical models are warranted to further define alterations in the B cell responses throughout *Ot* infection and long-lived humoral immunity after host recovery. Since other Gram-negative bacteria have been shown to selectively impair B cell activation [26], the use of low-virulence *Ot* strains, full spectral flow cytometry, and single-cell RNAseq will help understand strain-specific B cell responses, distinct B cell subpopulations, and dynamic transcriptomic signatures during scrub typhus progression.

Overall, this is the first study regarding B cell responses to acute *Ot* infection, demonstrating profound alterations at the tissue level (disorganized GC structures) and at the cellular/molecular level (impaired B cell activation and related pathways) during lethal scrub typhus in mice. Our findings provide valuable insights into the humoral immune response to *Ot* infection and open future research directions. A better understanding of B cells responses during acute and subclinical infections will provide vital knowledge that may better scrub typhus vaccine design.

## Acknowledgements

We would like to thank the UTMB Flow Cytometry and Cell Sorting Core Lab (Meredith Weglarz) the Optical Microscopy Core for our sample analyses, Joseph Thiriot for his help in mouse sample collection, and Dr. David Walker for providing the BSL-3 research facilities.

## Supporting Information

S1 Table. Primer sequences for qRT-PCR analysis used within the study.

**S1 Fig. Overview of differentially expressed genes identified by RNAseq during lethal *Ot* infection.** Mice were infected, as described in Figure 1; splenic B cells were purified from mock samples (n = 2) and at D4 (n =3) and D8 (n =3) for RNAseq analyses. Data were normalized via Relative Log Expression (RLE) using DESeq2 R library. Differentially expressed genes for from D4 and D8 samples identified by ROSALIND analysis, relative to mock, are shown in the A) bar graph and B-C) volcano plots.

**S2 Fig. Differential expression of B cell activation genes on D4 of lethal *Ot* infection.** Mice were infected, as described in Figure 1; splenic B cells were purified from mock samples (n = 2) and at D4 (n =3) and D8 (n =3) for RNAseq analyses. Data were normalized via Relative Log Expression (RLE) using DESeq2 R library. Differentially expressed genes for B cell activation (PANTHER and Gene Ontology Biological Process databases) from mock and D4 samples were identified by ROSALIND analysis and listed in the heatmap as subtracted normalized Log2 expression values.

**S3 Fig. Differential expression of BCR signaling genes on D8 of lethal *Ot* infection.** Mice were infected, as described in Figure 1; splenic B cells were purified from mock samples (n = 2) and at D4 (n =3) and D8 (n =3) for RNAseq analyses. Data were normalized via Relative Log Expression (RLE) using DESeq2 R library. Differentially expressed genes for BCR signaling (KEGG and Gene Ontology Biological Process databases) from mock and D8 samples were identified by ROSALIND analysis and listed in the heatmap as subtracted normalized Log2 expression values.

